# Mutualism in disguise? Isotopic evidence for nutrient transfer from a carnivorous pitcher plant to its insect prey

**DOI:** 10.1101/2025.10.16.682955

**Authors:** David W. Armitage, Asa Conover, Katharine M. SaundersM

**Affiliations:** Integrative Community Ecology Unit, Okinawa Institute of Science and Technology Graduate University, Onna, Okinawa 904-0495 Japan; Department of Integrative Biology, University of California, Berkeley, California 94720 USA

**Author notes:** Author roles*: DWA conceived the study. All authors contributed to data collection, reviewing, and editing the manuscript.

**Keywords:** Pitcher plant, *Darlingtonia*, predation, mutualism, stable isotopes, carnivorous plants

## Abstract

Carnivorous pitcher plants are regarded as exploiters of arthropod prey, attracting them with nectaries which are densely clustered near the slippery peristome, putting visiting arthropods at risk of becoming trapped. However, most pitcher visitors are not captured and may benefit from access to concentrated, nutrient-rich rewards. This raises the possibility that pitcher plants and their arthropod visitors engage in an indirect mutualism in which both insects and plants incur some cost but reap net nutritive benefits, yet evidence that potential prey species derive measurable nutritional benefits from pitcher plants is currently lacking. To address this knowledge gap, we measured levels of nitrogen isotopic enrichment of vespulid wasps residing in dense stands of the naturally ^15^N enriched pitcher plant *Darlingtonia californica* relative to those from adjacent forest patches. Wasps occupying pitcher fens had elevated *δ*^15^N values, suggesting they consume ^15^N-enriched nitrogen originating from *Darlingtonia* — either directly through nectar or indirectly via ^15^N-enriched arthropod proteins. These findings contribute support for the hypothesized nutritional mutualism between pitcher plants and local insect populations.

## Introduction

Carnivorous pitcher plants present a collection of visual (Joel et al., 1985; Moran, 1996), olfactory (Di Giusto et al., 2010), and nectar lures (Bennett and Ellison, 2009; Cipollini et al., 1994) to capture prey. However, most visitors to pitcher traps are not captured but instead appear to exploit the pitchers for food and shelter (Adlassnig et al., 2011; Lam and Tan, 2018). This leads to the prediction that carnivorous pitcher plants may support their local insect prey populations through resource provisioning, which recasts the traditionally ascribed predator-prey interaction between pitcher plants and insects as a mutualism with net benefits to populations of both the plant and its arthropod prey (Joel, 1988).

Maintenance of such a mutualism requires the reward to both plant and visitor to outweigh both the cost of visitor attraction by the plant and *per capita* mortality risk by the visitor. This may be the case for visitors to the pitcher plant *Darlingtonia californica* which only captured 1.27% of vespulid wasps visiting its leaves over hundreds of hours of visual surveys (Dixon et al., 2005). Similarly low *per capita* capture rates have been documented in pitchers of *Sarracenia* (Cresswell, 1991; *Gibson, 1983; Heard, 1998; Newell and Nastase, 1998), Nepenthes* (Bauer et al., 2009; *MacFarlane, 1893), and Heliamphora* (Jaffe et al., 1992).

Conversely, pitcher plants also incur costs when producing attractants. The cost of secreting nectar is evident in its timing and location: it is produced only during the first few months of a pitcher leaf’s yearlong lifespan and is concentrated almost entirely near the entrance of the trap (Cipollini et al., 1994). Like other plant nectars (Nicolson and Thornburg, 2007), pitcher plant nectar contains a variety of amino acids (Dress et al., 1997), which represent a costly use of prey-derived nitrogen (N) in an N-limited habitat. Given the relatively low rates of prey capture, and low photosynthetic and nutrient use efficiencies documented in pitcher plants (Ellison and Farnsworth, 2005; Gilbert et al., 2025), their costly production of nectar appears to outweigh the consequences of not doing so.

If the production of nectar by pitcher leaves directly or indirectly contributes to the nutrient budgets of local prey insects it could create a feedback loop in which an energetically costly attractant also supplements prey populations (Joel, 1988). This possibility is supported by the experimental finding that non-prey insect abundance is positively correlated with pitcher density at the scale of individual plants (Gibson, 1983). Further evidence for this feedback would include clear signs that pitcher plants provide meaningful dietary supplementation to prey populations. This could occur either directly via nectivory/folivory, or indirectly via scavenging of drowned prey or simply predation on other insects using the pitcher leaves. While various methods exist for characterizing insect diets, stable isotope analysis offers a particularly powerful and practical approach for quantifying the relative contributions of carnivorous and non-carnivorous plant sources to arthropod biomass.

Stable isotopes of nitrogen can be used to trace dietary sources ecological communities (Fry, 2006), with δ^15^N values typically increasing 2–4‰ per trophic level (Post, 2002). Carnivorous pitcher plants, which obtain most of their nitrogen from captured arthropod herbivores and predators, are expected to exhibit *δ*^15^N values elevated between 2 and 8‰ relative to neighboring noncarnivorous plants (Givnish and Shiba, 2022; Schulze et al., 1997). Therefore, arthropods which feed either directly on pitcher plants (via nectar, roots, or leaf tissue), or on other arthropod populations that are supplemented by pitcher plant nitrogen, are expected to show *δ*^15^N enrichment relative to individuals of the same species occupying neighboring sites which lack carnivorous plants. If pitcher feeding represents a substantial resource base for a species, then this trophic enrichment is anticipated across a sample of the target species’ population within a local patch.

We tested this hypothesis by comparing the natural abundance 15N ratios of *Vespula* wasps collected either within sites dominated by the pitcher plant *Darlingtonia californica* or from nearby forest habitats lacking any carnivorous plants. Significant enrichment of *δ*^15^N in wasps from pitcher fens is interpreted as evidence that their nitrogen budgets are partially supplemented by pitcher plants, supporting the hypothesis that pitcher plants and their insect visitors may engage in a form of reciprocal nutrient provisioning.

## Methods

### Natural history of *Darlingtonia*

*Darlingtonia californica* Torr. (1853) (Sarraceniaceae) is native to Northern California and Oregon, where it inhabits spring-fed fens, streams, and ponds flowing over serpentine outcrops from sea level to 2100 m. It uses both chemical and visual lures to catch prey, including dense nectaries near the pitcher opening, translucent fenestrae, and a combination of slippery waxes, trichomes, and fluid to retain and drown prey (Adams and Smith, 1977; Armitage, 2016a,b). Its prey are typical for the family and includes ants, flies, wasps, beetles, and spiders (Cresswell, 1991; Heard, 1998; **?**). Frequent visitors to the pitcher leaf’s nectaries include social wasps, butterflies, and ants (**Figure 1A-C, Supplemental Video S1**).

**Figure 1.**
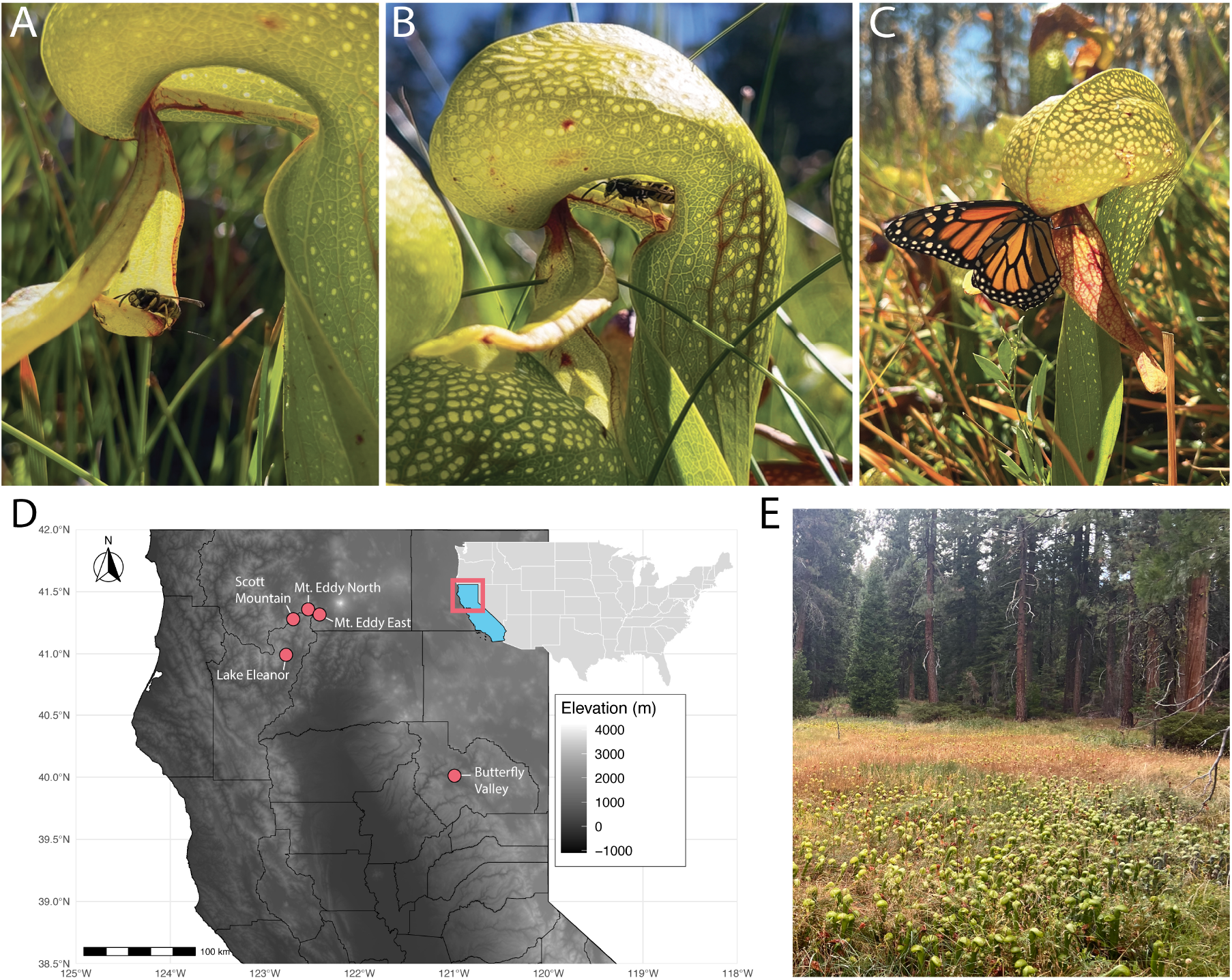
Vespulid wasps (*A, B*) and a butterfly (*C*) feeding on nectar from the trap entrances of *Darlingtonia californica* pitcher leaves. (*D*) Map of wasp collection sites in California, USA. (*E*) Typical montane fen containing *Darlingtonia californica* in the study area. Photo credits: A. Conover.

### Sample collection and processing

Sampling was conducted during August 2022 (Plumas County, CA) and October 2024 (Trinity County, CA and Siskiyou County, CA). We identified five accessible montane fens in which *Darlingtonia californica* plants grow at high abundances relative to sites across the range (**Figure 1D**). These sites contained thousands of individual plants spread evenly over areas between 0.5 and 1.5 hectares (**Figure 1E**). We also identified control sites approximately 1 km away from each *Darlingtonia* fen in the semi-open forest typical of the area. Insects captured at these sites were assumed to be unaffected by *Darlingtonia* nearby, as *Vespula* wasps typically forage within 300 meters of their nests to forage on other arthropods, carrion, fruit, and nectar (Akre et al., 1975).

At each of these sites, a single yellowjacket trap (Rescue! co., ltd) was hung on a tree branch in a relatively open area. This trap uses color and chemical attractants to quickly capture and kill vespulid wasps. Traps were deployed for between 1 and 5 days before being recovered. Identification was carried out and it was determined that the wasps trapped in the Trinity Co. and Siskiyou Co. traps were almost entirely comprised of *Vespula pensylvanica*, while wasps trapped in Plumas Co. were entirely *Vespula atropilosa*. On the day of collection, wasps were moved into silica beads for drying and shipping. Once dry, twelve individual wasps per trap were randomly removed from the vials and pulverized. This sample size was chosen to balance cost against statistical robustness and site number. These powdered wasps were then weighed on an ultra-microbalance and placed into tin capsules for isotope analysis. Alongside these samples, we submitted powdered samples from *Darlingtonia* leaves as well as other non-carnivorous plant taxa occurring in the pitcher fens and neighboring forest sites to evaluate levels of isotopic Nitrogen enrichment in *Darlingtonia* relative to the plant baseline. Isotope analysis was carried out at the University of California Davis Stable Isotope Facility on a Carlo Erba NC2500 elemental analyzer (Thermo Fisher Scientific, Inc.) interfaced to a Sercon 20-22 isotope ratio mass spectrometer (Sercon Ltd., Cheshire, UK). Isotope masses were transformed to *δ* notation as

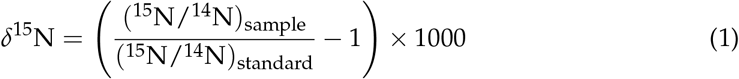

where the numerator is the ratio of heavy to light isotopic mass of the sample and the denominator is the equivalent ratio of the atmospheric ^15^N standard (*≈* 3.677 *×* 10^*−*3^).

### Statistical analysis

The *δ*^15^N distributions of each source material were compared using empirical and fitted theoretical cumulative distribution functions. We then applied estimation statistics to calculate the bootstrapped mean difference in *δ*^15^N values between wasps from pitcher fens and those from forest sites (Ho et al., 2019). Gardner-Altman plots showing bootstrapped mean distances between groups were constructed using the R package dabestr (Ho et al., 2019).

While estimation statistics are well-suited for comparing mean differences, they do not account for site-specific variation in nitrogen sources unrelated to the presence or absence of *Darlingtonia*. To address this and evaluate the probabilistic evidence for *Darlingtonia*-mediated *δ*^15^N enrichment in wasps, we fit a linear mixed-effects model to the *δ*^15^N data. Habitat type (fen vs. non-fen control) was included as a fixed effect, with site ID modeled as a random intercept.

Modeling was implemented in the R package brms (Bü rkner, 2017), an R frontend for the Stan language (Carpenter et al., 2017). Models assumed a Gaussian error structure, an improper flat prior for pitcher fen effects, and weakly informative *t*-distributed priors for other effects (to prevent extreme values or ensure positive sign). Six MCMC chains were run for 25,000 post–burn-in iterations, thinning every 20^th^ iteration for 7,500 total posterior draws. Mixing and convergence was verified using 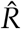 and effective sample sizes (*N*_*e f f*_) (Vehtari et al., 2021). We then calculated the one-sided posterior probability that the pitcher fen effect was greater than zero, indicating isotopic enrichment of wasps in pitcher fens compared to control forest.

We also investigated the influence of individual data points on model fit using Pareto smoothed importance sampling (PSIS), a method for identifying influential observations in leave-one-out (LOO) cross-validation (Vehtari et al., 2024). One data point had Pareto *k* values *>* 0.7, indicative of a high-leverage outlier, but removing it and recalculating the bootstrapped mean differences and model posterior distributions resulted in nearly identical point estimates and error distributions. Alternative model specifications to the Gaussian random-intercepts model were also considered, including a random intercepts and slopes model, as well as Student-*t* distributions for both random effects models. However, loss of predictive density from PSIS-LOO did not support these more complex models over the Gaussian random-intercepts model (Δ_*elpd*_ *> −*2 for all models). Posterior predictive checks verified this model had an acceptable fit to the data.

## Results

Exploratory analysis of our data hinted at a potential isotopic separation of background fen vegetation and *Darlingtonia* plants, as well as as a much smaller separation between forest and pitcher fen wasps (**Figure 2A**). This was confirmed by our nonparametric bootstrapping support for the significant *δ*^15^N enrichment of *Darlingtonia* tissue relative to to that of background plants (mean difference = 6.8‰, *CI*_95%_ = [5.5, 8.9]), corresponding to an approximate increase of between two and three trophic levels. Similarly, the mean difference between pitcher fen wasps and background forest wasps was also greater than 0 (mean difference = 0.41‰, *CI*_95%_ = [0.12, 0.75](**Figure 2B**), but did not indicate a shift in trophic level.

**Figure 2.**
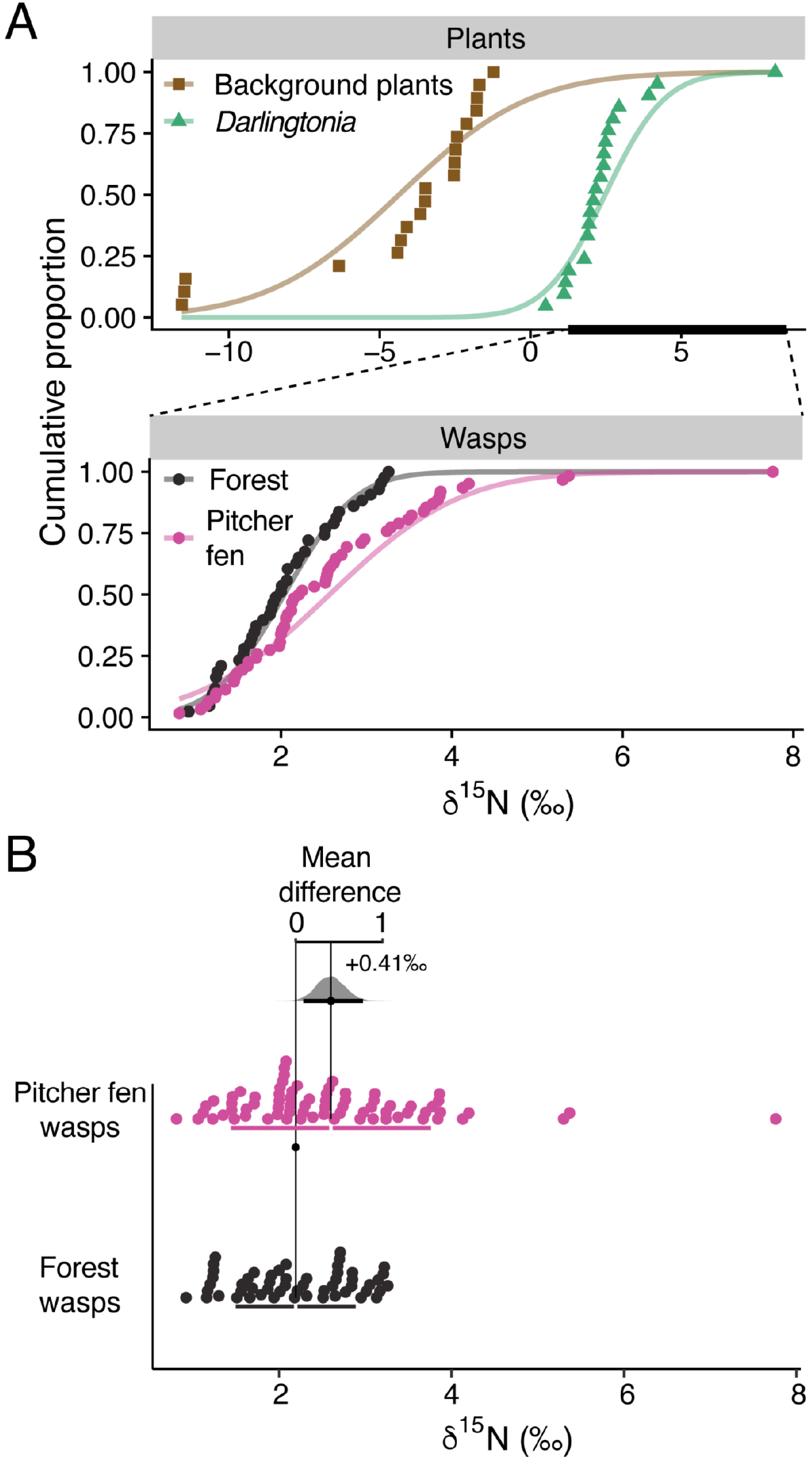
(*A*) Cumulative distributions of *δ*^15^N (relative to air) for individual samples. Plants are divided into background species versus carnivorous *Darlingtonia*; wasps are grouped by trap location (pitcher fens vs. forest controls). Points represent individual samples; lines indicate fitted theoretical distributions. (*B*) Gardner–Altman plot comparing wasp *δ*^15^N between fen and forest sites. Lower panel shows individual values with group means; upper panel shows the bootstrap distribution of the mean difference. The mean difference was 0.41 ‰ (95% CI [0.115, 0.756]).

From the posterior distribution of our Bayesian linear mixed-effects model, we tested the directional hypothesis that wasps captured in pitcher fens exhibit elevated *δ*^15^N values compared to those from forest reference sites. This prediction was strongly supported, with a one-sided posterior probability of 0.98 that the effect was greater than zero (mean effect = 0.35‰, *CI*_95%_ =]0.02, 0.58], evidence ratio = 55.4) (**Supplemental Figure S1**), even after accounting for random, location-specific variation in baseline *δ*^15^N values. However, the proportion of variance explained by the model was relatively modest (Bayesian *R*^2^ = 0.21 *±* 0.05), indicating that a substantial amount of variation remains unexplained.

## Discussion

The hypothesis that carnivorous pitcher plants may serve as nutrient sources for local arthropod populations — rather than purely as predators — is supported by our finding that yellowjacket wasps collected in dense stands of *D. californica* pitchers exhibit a modest, yet significant enrichment in *δ*^15^N relative to conspecifics from neighboring sites that lack carnivorous plants. This isotopic signal is consistent with nitrogen assimilation from pitcher plants, which exhibit naturally enriched ^15^N levels indicative of a two- to three-trophic level increase. Similar *δ*^15^N enrichment has also been documented in the inquiline larval insects and adult ants associated with the pitchers of *Nepenthes bicalcarata*, relative to the isotopic composition of the plant’s prey items (Bazile et al., 2012; Scharmann et al., 2013). It remains to be determined whether this nitrogen is acquired directly via nectar feeding or indirectly via consumption of prey populations supported by pitcher-derived nutrients.

One potential explanation for the only moderate effect size (Hedge’s *g* = 0.42) observed in the pitcher fen wasp enrichment is that pitcher leaf nectar (like most plant nectars) contains little nitrogen and therefore contributes minimally to wasps’ nitrogen content. However, the marked enrichment of pitcher *δ*^15^N relative to surrounding vegetation suggests that trophic ^15^N enrichment may still occur, even if wasps assimilate only small amounts of nitrogen from nectar. Social wasps acquire most of their nitrogen during the larval stage, when building tissues such as muscle and exoskeleton. In contrast, adults are no longer growing and have limited nitrogen needs, mainly using it for short-lived enzymes and proteins, the ^15^N enrichment of which would be diluted by the aforementioned tissues. In many nectivorous animals amino acids from nectar can provide a substantial subsidy to the lifetime N requirement (Nicolson, 2007).

An alternative pathway for ^15^N enrichment is through the scavenged animal proteins — themselves enriched from pitcher tissues — fed to larvae by adult workers. As no larvae or their arthropod prey were collected in our surveys, both pathways remain plausible and may operate in tandem. In either case, however, the pitcher plants form the basis of this nutrient loop, channeling prey-derived nitrogen back into the insect populations they have evolved to attract. Such reciprocal flows reveal a more subtle ecological role for these carnivorous plants within their local food webs. Despite the overall enrichment of wasps in pitcher fens, this trend was evident at only three of the five sites—Mount Eddy East and North showed no detectable difference between fen and forest populations. This inconsistency suggests that yellowjacket populations may vary in their use of pitcher plants or in the trophic niches they occupy (*sensu* Torniainen and Komonen (2021)). Future work could test this directly through standardized surveys of wasp abundance and diet composition within and outside pitcher fens.

Even so, such variation would not preclude the existence of a more diffuse interaction in which pitcher plants enrich the local prey base and indirectly elevate the trophic status of wasps foraging in these habitats. For instance, wasps in montane fens may feed more frequently on predatory arthropods (e.g., spiders), whereas those in adjacent forests may rely on herbivorous prey such as beetles. In this case, the observed isotopic enrichment could reflect a form of trophic upgrading mediated by the fen food web itself, with *Darlingtonia* acting as a foundation species that enhances food web complexity. However, because we were unable to sample paired fen sites lacking *Darlingtonia* — which are rare in the study region — the specific contribution of the plant, independent of fen habitat effects, remains unresolved.

It is important to note, however, that evidence of dietary enrichment alone does not fully establish the net fitness benefit for visiting wasps required to concretely reclassify this interaction as mutualistic, but rather acts as a line of evidence supporting this interaction. Our isotope data demonstrate that wasps may assimilate nitrogen derived from *Darlingtonia*, but they do not reveal whether this subsidy outweighs the potential opportunity costs or risks associated with foraging at pitcher leaves. From the wasps’ perspective, then, the interaction could remain neutral or even costly. Thus, while our findings are consistent with a nutritional pathway linking pitcher plants and insect visitors, they fall short of demonstrating mutualism in the strict sense of reciprocal fitness benefits. However, the plant’s benefits from insect capture are evident, as reflected in its evolved structural adaptations that promote insect visitation and nutrient exchange, and reduced photosynthetic rates, vegetative growth, and seed production in experimental prey removal experiments (Farnsworth and Ellison, 2008; Ne’eman et al., 2006).

In conclusion, our stable isotope analyses indicate that yellowjacket wasps foraging in *Darlingtonia* fens incorporate pitcher-derived nitrogen into their diets, providing evidence for a reciprocal nutrient exchange between carnivorous plants and local insect populations. This finding challenges the traditional view of pitcher plants as strict predators, instead positioning them within a spectrum of plant–animal interactions that includes mutualistic nutrient exchange. Looking forward, studies combining direct behavioral observations, manipulative pitcher-removal experiments, and additional pitcher plant and arthropod species will be essential for identifying the explicit pathways of nitrogen transfer and for reconstructing the food webs of pitcher plant–dominated fens and bogs. Such work will help clarify the role of these unique plants as foundational species that support communities of animals markedly distinct from those in the surrounding habitats.

## Acknowledgments

Permits for sample collection were granted by the US Forest Service. Funding was through a subsidy from the Japanese Cabinet Office to OIST.

---

**Figure S1.**
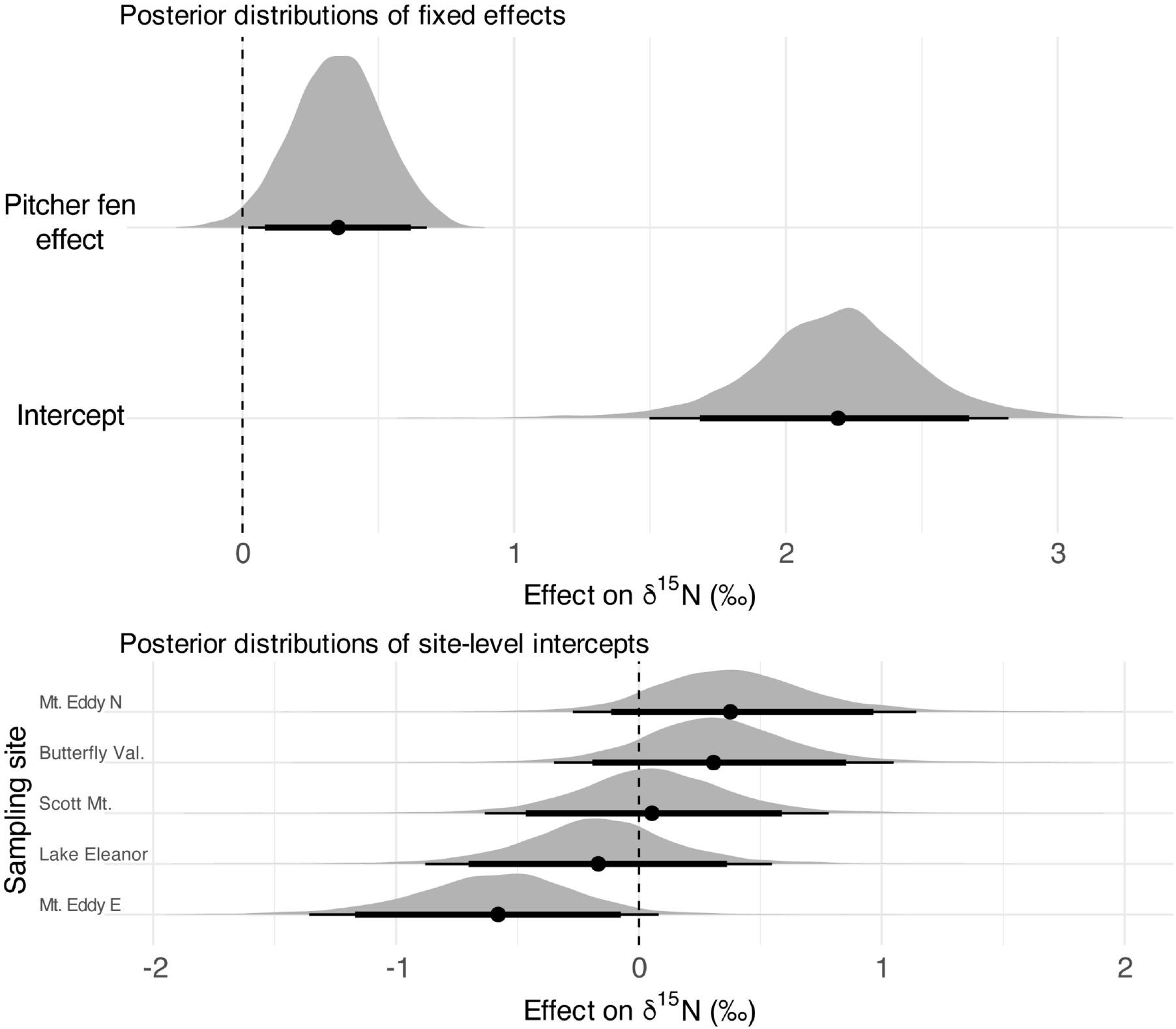
Posterior distributions of parameter estimates from a Bayesian linear mixed-effects model predicting *δ*^15^N values in wasps. The top panel shows the estimated effect of proximity to carnivorous pitcher plants (the pitcher fen effect) on *δ*^15^N, along with the global intercept. The bottom panel shows site-specific random intercept deviations in baseline *δ*^15^N values. Horizontal bars represent 89% and 95% credible intervals, and shaded densities reflect the full posterior distributions. Here, 98% of the posterior support for the pitcher fen effect lies above zero.

